# The *Wingless* Planar Cell Polarity pathway is essential for optimal activity-dependent synaptic plasticity

**DOI:** 10.1101/2023.12.01.569625

**Authors:** Carihann Dominicci-Cotto, Mariam Vazquez, Bruno Marie

## Abstract

From fly to man, the Wingless (Wg) / Wnt signaling molecule is essential for both the stability and plasticity of the nervous system. The *Drosophila* neuromuscular junction (NMJ) has proven to be a useful system for deciphering the role of Wg in directing activity-dependent synaptic plasticity, which, in the motoneuron, has been shown to be dependent on both the canonical and the noncanonical calcium Wg pathways. Here we show that the noncanonical planar cell polarity (PCP) pathway is an essential component of the Wg signaling system controlling plasticity at the motoneuron synapse. We present evidence that disturbing the PCP pathway leads to a perturbation in activity-dependent synaptic plasticity (ADSP). We first show that a PCP-specific allele of *dishevelled* (*dsh*) affects the *de novo* synaptic structures produced during ADSP. We then show that the Rho GTPases downstream of Dsh in the PCP pathway are also involved in regulating the morphological changes that take place after repeated stimulation. Finally, we show that Jun kinase is essential for this phenomenon, whereas we found no indication of the involvement of the transcription factor complex AP1 (Jun/Fos). This work shows the involvement of the neuronal PCP signaling pathway in supporting ADSP. Because we find that AP1 mutants can perform ADSP adequately, we hypothesize that, upon Wg activation, the Rho GTPases and Jun kinase are involved locally at the synapse, in instructing cytoskeletal dynamics responsible for the appearance of the morphological changes occurring during ADSP.

## Introduction

Synapses are the site of plastic events that shape their morphological and electrophysiological properties depending on their experiences (Harris & Littleton, 2015; Ho et al., 2011; Holtmaat & Svoboda, 2009; Pereda, 2014; Shepherd, 2004). One such plastic event is activity-dependent synaptic plasticity (ADSP), by which the synaptic efficacy can be increased or decreased, reflecting previous excitatory or inhibitory stimuli (Castillo, 2012; Lüscher & Malenka, 2012). This plasticity is thought to be the cellular correlate of learning and memory, and it is essential to improve our understanding of the molecules and molecular signals underlying this phenomenon. The secreted molecule Wg/Wnt regulates ADSP (Budnik & Salinas, 2011; Rosso et al., 2013). Indeed, activity-dependent mechanisms induce Wnt release that, in turn, induces plasticity signaling in hippocampal neurons (Chen et al., 2006; Tabatadze et al., 2014). This increase in Wnt secretion provokes structural alterations in dendritic arborizations (Ferrari et al., 2018; Wayman et al., 2006; Yu & Malenka, 2003) and spines (Ciani et al., 2011; McLeod et al., 2018; Tabatadze et al., 2014), promoting changes in synaptic strength and plasticity. *Drosophila* served as a model of choice to explore the instrumental role of Wg in directing ADSP (Bai & Suzuki, 2020; Budnik & Salinas, 2011). ADSP can be elicited at the NMJ when the preparation is submitted to a repeated stimulation protocol (Ataman et al., 2008; Maldonado-Díaz et al., 2021; Piccioli & Littleton, 2014; Vasin et al., 2014, 2019), similar to the one used on hippocampal neurons to elicit LTP (Wu et al., 2001). The changes associated with the NMJ experiencing ADSP are both electrophysiological and structural (Alicea et al., 2017; Ataman et al., 2008; Maldonado-Díaz et al., 2021). The structural changes consist of the formation of *de-novo* synaptic structures named “ghost boutons” (Fig. 1A), and the Wg canonical pathway is essential for this plasticity (Ataman et al., 2008). Repeated stimulation of the NMJ promotes an increase in synaptic Wg that leads to rapid structural and electrophysiological changes (Ataman et al., 2008). In fact, Wg signaling regulates these changes bidirectionally; post-synaptically through nuclear import of its receptor, and pre-synaptically by the inhibition of GSK3ß/Sgg (Ataman et al., 2008). In addition, there is evidence that the calcium pathway is involved in this plasticity. In this pathway, an increase in intracellular calcium activates CaMKII and the transcription factor NFAT (Koles & Budnik, 2012). The neuronal expression of NFAT subunit A at the NMJ prevents the formation of ghost boutons upon stimulation (Freeman et al., 2011). In addition, synaptic CaMKII protein expression increases upon stimulation which correlates with *de novo* bouton formation (Nesler et al., 2016). However, to date the third pathway that responds to the Wg/Wnt signal, the planar cell polarity (PCP) pathway, has not yet been assessed for its role in regulating ADSP.

**Figure 1:**
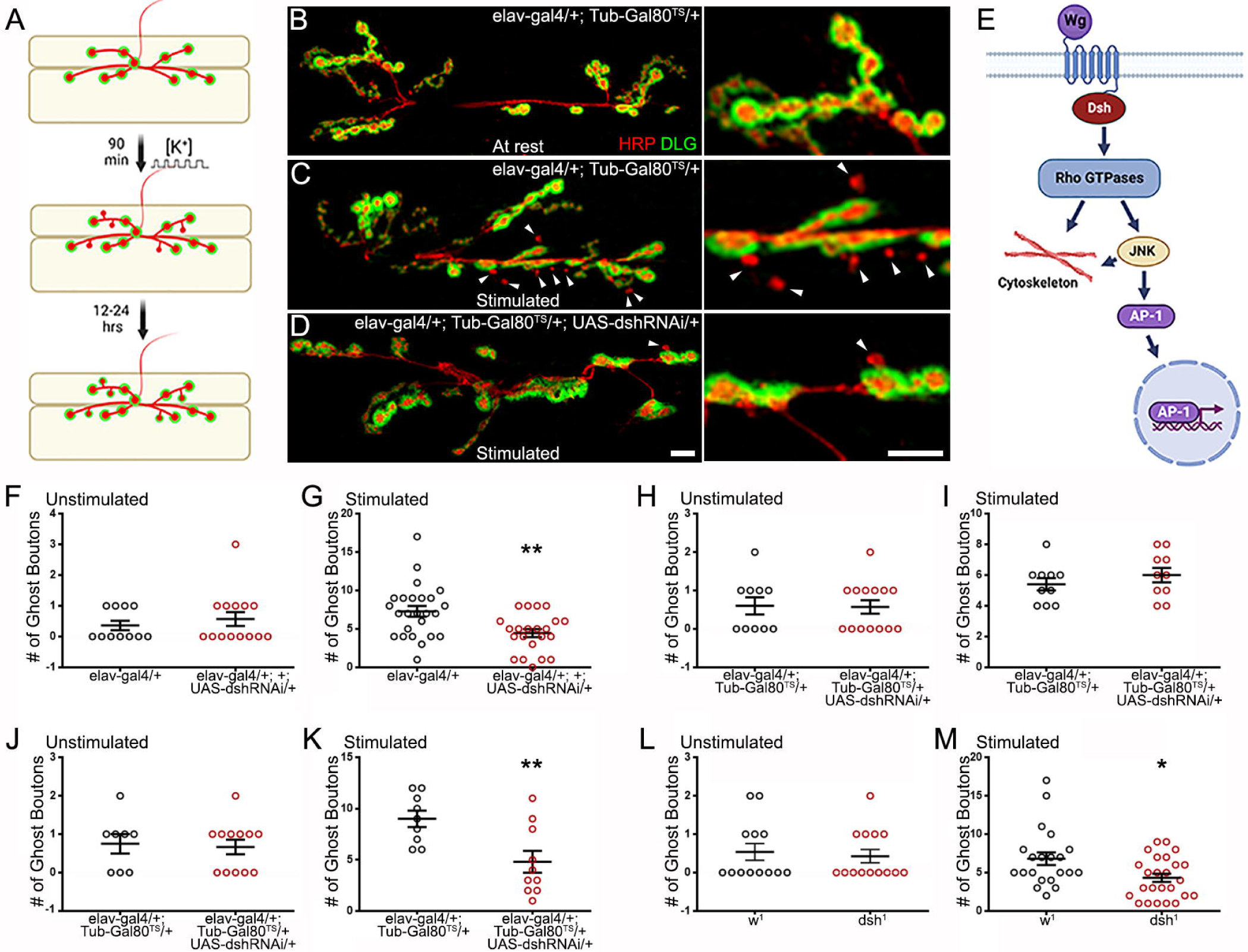
Dishevelled and its planar cell polarity domain regulate ADSP. **(A)** Schematic diagram representing a NMJ labeled with pre-synaptic (red) and post synaptic (green) markers. Upon stimulation, *de novo* synaptic boutons called “ghost boutons” are identifiable because they only show the pre-synaptic marker HRP. These structures, with time, will acquire post-synaptic differentiation. **(B)** Representative confocal photograph of an unstimulated control synapse showing HRP (presynaptic marker) and DLG (postsynaptic marker) immunoreactivity. **(C)** Representative confocal photograph of a stimulated control synapse. Arrowheads point to ghost boutons. **(D)** Representative confocal photograph of a stimulated synapse expressing *dsh* RNAi. **(E)** Schematic diagram illustrating the components of the Wg planar cell polarity pathway. **(F-M)** Quantification of the ghost bouton number in **(F)** unstimulated and **(G)** stimulated control and preparations expressing *dsh* RNAi constitutively; **(H)** unstimulated and **(I)** stimulated control and preparation carrying the suppressed conditional expression system at 18°C; **(J)** unstimulated and **(K)** stimulated control and preparation expressing *dsh* RNAi for 48 hours prior to the stimulation (activated expression system at 29°C); **(L)** unstimulated and **(M)** stimulated control and dsh^1^ mutant preparations. Scale bar: 10 µm. Two-tailed, unpaired t-tests: *p < 0.05; **p < 0.01. Data are shown as scatter plots and means ± SEM.

Here we use the *Drosophila* NMJ model to ask whether the Wg PCP pathway (Fig. 1E) is involved in ADSP. We use the appearance of *de novo* synaptic structures, ghost boutons, to quantify the extent of these changes. Because of their rapid development (90 min following stimulation), ghost boutons represent early pre-synaptic structures that are devoid of post-synaptic differentiation (Fig. 1A). This characteristic has been widely used to easily identify ghost boutons at the NMJ, in order to quantify ADSP (Alicea et al., 2017; Ataman et al., 2008; Freeman et al., 2011; Fuentes-Medel et al., 2009; Maldonado-Díaz et al., 2021; Nesler et al., 2013, 2016; Piccioli & Littleton, 2014; Vasin et al., 2014, 2019). Our present work investigates the role of the PCP pathway in the production of these morphological synaptic changes. We find that several pathway members are involved in this process. Indeed, we first show that a PCP-specific *dsh* allele (Perrimon & Mahowald, 1987) is necessary for optimal ADSP. We then show that activity of the Rho, Rac1, and Cdc42 GTPases can also influence ADSP. We then investigate the Jun kinase (JNK), an effector of these GTPases (Coso et al., 1995; Rosso et al., 2005) and show that its loss of function is detrimental to ADSP; however, the transcription factor AP1 (Fos/Jun), a target of JNK (Etter et al., 2005; Sanyal et al., 2002), is not necessary for this process. In most of our experiments, we use an inducible RNA expression system to selectively down- or up-regulate the activity of the members of the PCP pathway pre-synaptically only (post-synaptic expression is unchanged) and late in development (after the establishment of the NMJ). This allows us to conclude that the effects we observed are due to a change in the PCP pathway activity within the motoneuron and not a consequence of developmental perturbation. On the contrary, we argue that it reflects a need for the PCP pathway within the motoneuron to achieve optimal ADSP.

## Material and Methods

### Genetics

For experiments using the *elav^c155^-gal4* insertion on the X chromosome, only females were used (in order to consider heterozygote animals). In the other experimental procedures, animals from both sexes were used. Flies were reared on standard fly food at 18°C for conditional experiments or at 25°C in all other cases. Our methods are consistent with standard husbandry and care. Our laboratory follows ethical practices while carrying out experiments as well as the cataloguing of *Drosophila*. The fly strains used in this study include *dsh^1^* [Bloomington Drosophila stock center (BDSC), stock # 5298], and *w^1^* (BDSC, stock # 145) as a control. We used the Gal4/UAS system (Brand & Perrimon, 1993) to express dominant-negative or constitutively active genes in neurons. For this, we used the pan-neuronal driver *elav^c155^-gal4* (BDSC, stock # 458) whose expression starts early in embryogenesis (after neuroblast mitosis) in combination with *UAS-dsh-RNAi* (BDSC, stock #31306), *UAS-rac1^N17^*(BDSC, stock #6292), *UAS-rac1^V12^* (BDSC, stock # 6291), *UAS-rho1^N19^*(BDSC, stock # 58818), *UAS-rho1^V12^* (BDSC, stock # 58817), *UAS-cdc42^N17^*(BDSC, stock # 6288), *UAS-cdc42^V12^* (BDSC, stock # 4854), *Tub-Gal80^TS^*(BDSC, stock #7108), *UAS-jnk^DN^* (BDSC, stock # 9311), *UAS-fos^DN^*(BDSC, stock #7215) and *UAS-jun^DN^* (BDSC, stock #7218). The controls for these experiments were the driver elav^c155^-gal4 / +. For the conditional experiments, we made a fly containing the *elav^c155^-gal4* on the first chromosome and *Tub-Gal80^TS^* on the second chromosome. This fly expresses the thermosensitive inhibitor Gal80^TS^ (McGuire et al., 2004) under the control of by ubiquitous *tubulin* (*Tub*) promoter. Larvae for the conditional experiments were reared at 18°C, where Gal80^TS^ inhibits protein expression by binding to the Gal4 activator. Then, 48 hours before performing the stimulation protocol, we shifted them to 29°C, inhibiting Gal80^TS^ and therefore allowing the expression of specific proteins activated by Gal4.

### Stimulation protocol

We dissected the body wall muscles from third instar larvae in hemolymph-like HL3 saline (70 mM NaCl, 10 mM NaHCO3, 115 mM sucrose, 5 mM trehalose, 5 mM HEPES, 10 mM MgCl2), leaving unharmed the CNS and the peripheral nerves innervating the body wall muscles. The protocol was carried out on partially intact and non-stretched preparations. The stimulation protocol that we used was adapted from Ataman et al., 2008 and is constituted of 5 stages that alternate stimulation and rest periods. In the first three stages the preparations were stimulated for 2 min followed by a 15 min rest period. The fourth stage is composed of a 4 min stimulation period followed by a 15 min rest while the fifth and last stage is composed of a 6 min stimulation period followed by a 14 min rest. The 5 stimulation periods were elicited using a potassium- and calcium-rich HL3 saline solution (90 mM KCl and 1.5 mM CaCl2), while during the five rest periods a HL3 saline containing 5 mM KCl and 0.1 mM CaCl_2_ was applied.

### Immunohistochemistry

After stimulation, the preparations were fixed for 15 min at room temperature in a solution of 4% paraformaldehyde in PBS before being incubated overnight at 4°C within a primary (anti-Dlg) antibody solution, (1:20; Budnik et al., 1996). The secondary antibody (1:300; Alexa Fluor 488-conjugated AffiniPure goat anti-mouse or anti-rabbit, Jackson ImmunoResearch) and Anti-Hrp (1:300; Cy3-conjugated AffiniPure goat anti-horseradish peroxidase, Jackson ImmunoResearch; Jan & Jan, 1982) were applied for 1 hr at room temperature as previously described (Alicea et al., 2017; Maldonado-Díaz et al., 2021).

### Quantification of ghost boutons

We treated control preparations in parallel with each experimental genotype to account for the potential variation in our experimental manipulations. The control preparations were designed to contain the experimental preparations’ genetic background (*w^1^* for *dsh^1^*or the corresponding *gal4* driver for the different expression systems used). Control and experimental preparations were also reared at the same temperature and submitted to the same shift (18°C to 29°C) when applicable. We identified a ghost bouton as a bouton that had a positive immunoreactivity to anti-HRP and negative immunoreactivity to anti-Dlg. In all the experiments, ghost boutons were counted on muscle 6/7 NMJs, in A3 segments of third instar larvae. To carry out these observations, we used a Nikon Eclipse 80i microscope at a magnification of 400X.

### Statistical treatment

We first identified any outlier using the ROUT method with a Q = 0.1%. Then, we assessed whether the data fitted a normal distribution. Using the Shapiro-Wilk normality test, we classified the data as parametric (p > 0.0001) or nonparametric (p < 0.0001). In cases, when the data were nonparametric, we ran a Kruskal–Wallis test with a post hoc Dunn’s multiple comparison test. For cases of parametric statistics, we ran an ANOVA test—when there are three or more data sets. We applied a post hoc Dunnett’s correction test when multiple comparisons are carried out against a control value. When only two data sets were compared, we performed two-tailed, unpaired t-tests. GraphPad Prism 6 was used to apply these statistical treatments.

## Results

### Disheveled and its PCP-specific domain relay the signal necessary for ADSP

Previous studies showed that the secreted molecule Wg and its pre-synaptic receptor Fz2 controlled ADSP at the *Drosophila* NMJ (Alicea et al., 2017; Ataman et al., 2008). Because Disheveled (Dsh) is the cytoplasmic molecule making the link between the Wg receptor and its different effectors, it is often called the Wg hub (Gao & Chen, 2010). We asked whether Dsh loss of function could reduce ADSP and phenocopy the defects in ADSP observed in the *wg* and *fz2* loss of function backgrounds (Alicea et al., 2017; Ataman et al., 2008). We used a well-described method to repeatedly stimulate the NMJ preparation in a way reminiscent of the stimulation protocol used on hippocampal neurons in culture (Wu et al., 2001). In these conditions, after a 90-min patterned repeated stimulation protocol, we observe the appearance of ghost boutons (see schematic diagram Fig. 1A; Ataman et al., 2008) that show the neuronal membrane-specific marker revealed by the anti-HRP immunoreactivity (Jan & Jan, 1982), without the apposition of the post-synaptic marker Dlg (Disc large, the membrane-associated guanylate kinase homolog; Budnik et al., 1996; Lahey et al., 1994). These *de-novo* synaptic structures are noticeable when comparing unstimulated control synapses (Fig. 1B) to stimulated control synapses (Fig 1C; arrowheads point to ghost boutons) and are easily quantifiable (Fig 1F-M).

Throughout this manuscript, we paid attention to the status of the synapse at rest (without stimulation) to ensure that there was no significant difference in the mean synaptic ghost bouton number between the studied genetic backgrounds. This way, we can infer that any observed differences after stimulation reflect a change in the synaptic answer to the activity-dependent process. In fact, unstimulated preparations across all the genotypes studied (Fig 1 F, H, J, L) show no differences. We later ensured that our stimulation protocol was efficient in eliciting ghost boutons. Here, our unstimulated control preparations showed a mean of 0.36 ± 0.15 ghost boutons per synapse (Fig. 1F, black circle, n = 11), and our stimulated controls showed 7.28 ± 0.69 ghost boutons (Fig 1G, black circle, n= 25), indicating an increase in ghost boutons comparable to previously published work (Alicea et al., 2017; Ataman et al., 2008; Freeman et al., 2011; Lee et al., 2017; Maldonado-Díaz et al., 2021; Nesler et al., 2013, 2016; Piccioli & Littleton, 2014; Vasin et al., 2014, 2019).

We then asked whether Dsh was necessary to perform ADSP. Because of the importance of Dsh in transducing the Wg signal, its null alleles (for example *dsh^X788^*and *dsh^nYn234Y^*) are embryonic lethal and therefore do not allow the study of the larval NMJ (Klingensmith et al., 1994; Perrimon & Mahowald, 1987). To create a viable loss of function condition for *dsh*, we used RNA interference (RNAi; Piccin et al., 2001) and expressed it using the *elav-Gal4* post-mitotic neuronal driver (Robinow & White, 1991). When these preparations were submitted to the patterned repeated stimulation protocol, they showed fewer ghost boutons (Fig 1.G, red circles). The mean ghost bouton number in these preparations was 4.46 ± 2.48, a value less than the control value (p = 0.0026; n = 22), suggesting that Dsh is indeed involved in ADSP.

Although we used the postmitotic driver *elav-Gal4* to initiate *dsh* loss of function, its expression starts early in development (end of embryogenesis). Therefore, the phenotype we observe could be caused by a detrimental event during synapse formation due to a lack of Dsh activity. To rule out this possibility, we first examined the NMJ and did not notice any phenotype affecting axon routing or synaptic growth. Secondly, we designed an inducible expression system to affect Dsh activity. This system involves the expression of 3 different transgenes within the studied animal: the *elav-Gal4* driver, the effector *UAS-dsh-RNAi*, and the inhibitor *Tub-Gal80^TS^*. In this system, the temperature-sensitive (TS) Gal80^TS^ protein is expressed ubiquitously (under the control of the tubulin promoter) and able to repress Gal4 at permissive temperature (18°C; McGuire et al., 2004). Indeed, at this temperature, the expression of *dsh* RNAi is shut off and we observe no difference between the controls and the animals containing *dsh* RNAi in both non-stimulated (Fig. 1H) and stimulated (Fig. 1I) preparations. At the restrictive temperature (29°C), the temperature-sensitive mutation carried by Gal80^TS^ is revealed, producing a non-functional Gal80, which leads to the de-repression of the system, and the expression of *dsh* RNAi. In this condition, we can observe a loss of function phenotype when we elicit ADSP. When we derepressed the Gal4/UAS system for 48 hours before performing the stimulus protocol, we did not observe any difference at rest between controls and dsh RNAi loss of function (the mean value of controls is 0.75 ± 0.25, n = 8, while it is 0.67 ± 0.19, n = 12 for dsh RNAi expressing animals; Fig. 1B and 1J). After repeated patterned stimulations, we did observe differences between controls and preparations containing the *dsh* RNAi construct. While the mean value of ghost boutons is 9 ± 0.8 for controls (n = 9; Fig. 1C and K), it is reduced in dsh RNAi animals (4.8 ± 1.07 ghost boutons; n=10; p = 0.007; Fig. 1D and K). This reduction, provoked by a loss of Dsh activity only 48 hours before applying the repeated stimulation protocol, is comparable to the reduction observed when we inhibit Dsh function since the end of embryogenesis (Fig. 1G). This suggests that the effect observed on ADSP is due to the requirement of Dsh function at the time of the stimulation challenge and not a consequence of perturbed development.

Having established the importance of Dsh in transducing the signal responsible for ADSP, we asked whether the planar cell polarity pathway (PCP) was involved in this phenomenon. To do so, we took advantage of the previously studied and isolated *dsh^1^* allele (Perrimon & Mahowald, 1987). This allele, often referred to as a PCP-specific allele, encodes a version of Dsh that selectively affects the transduction of the Wg signaling to the PCP pathway without altering the efficacy of the Wg signal toward the canonical or the calcium pathway (Axelrod et al., 1998; Boutros et al., 1998; Yanfeng et al., 2011). At rest, there is no difference between control and *dsh^1^* animals (the mean value of controls and *dsh^1^* animals are 0.54 ± 0.21, n = 13 and 0.55 ± 0.8, n = 14 respectively; Fig. 1L). Nevertheless, after repeated patterned stimulation, the mean number of ghost boutons appearing at the synapse is reduced in *dsh^1^* animals (4.32 ± 0.54, n = 25; red circles, Fig.1 M) when compared to control (6.81 ± 0.83, n = 21; p = 0.017; black circles, Fig.1 M). This result suggests that part of the Wg signal required to perform ADSP is transduced through the PCP pathway. To consolidate the idea that the Wg PCP pathway is an integral part of ADSP machinery, we decided to ask whether other members of this pathway (Fig.1 E) could affect this plasticity.

### Rho1 activity can repress the morphological changes associated with ADSP

Since Dsh transduces the Wg signal through several GTPases (Boutros et al., 1998; Fanto et al., 2000; Schlessinger et al., 2007), we decided to investigate the role of Rho1 in ADSP. Rho1 is a small GTPase that functions as a molecular switch cycling between an inactive GDP-bound form and an active GTP-bound form to mainly regulate the assembly and reorganization of the actin cytoskeleton controlling different cellular activities (Jaffe & Hall, 2005). In this way, Rho1 influences several biological processes such as embryogenesis, cell polarity, and cell division (Barrett et al., 1997; Hariharan et al., 1995; Olson et al., 1995;Strutt et al., 1997). Its role in neuronal development has also been studied; Rho activation can promote or inhibit neuronal growth cones as well as inhibit dendritic growth (Auer et al., 2011; Dickson, 2001; Luo, 2000). Reducing the activity of p190 RhoGAP, a negative regulator of Rho, provokes axonal guidance defects and decreases axon number in hippocampal cells (Brouns et al., 2001). In Drosophila mushroom bodies, the reduction of p190 RhoGAP promotes retraction of axonal branching, while its overexpression promotes axonal extension (Billuart et al., 2001). In addition, inhibition of neural activity in cortical neurons decreases active Rho levels while expression of constitutively active Rho increases axon branching by elimination and addition of branches (Ohnami et al., 2008).

Here we asked whether Rho1 was involved in ADSP at the NMJ. Because of Rho1’s multiple cellular functions its null mutations are lethal, so we therefore decided to express a dominant negative version (Rho1^DN^, see material and methods) and a constitutively active version (Rho1^CA^, see material and methods). We used the Gal4/Gal80^TS^ inducible system described previously to bypass the potential effects of these constructs’ expression on axon pathfinding and neuronal growth. We first checked that there were no notable differences regarding the number of ghost boutons in unstimulated preparations across all the genotypes considered at both permissive temperature (controls showed 1.21 ± 0.27 ghost boutons; n = 19; Rho1^DN^ 0.91 ± 0.28; n = 11; p = 0.57; Rho1^CA^ 0.64 ± 0.2; n = 14; p = 0.2 ; Fig. 2 D) and restrictive temperature (control showed 1.6 ± 0.29 ghost boutons; n = 22; Rho1^DN^ 1.14 ± 0.31; n = 14; p = 0.6; Rho1^CA^ 1.82 ± 0.28; n = 22; p = 0.97 ; Fig. 2 F). We also asked whether the inhibition of the Rho1^DN^ or Rho1^CA^ expression by Gal80^TS^ at 18°C was consistent with the observed results; we expected all the preparations to behave like controls if the expression was silenced efficiently. Indeed, the stimulated control preparations showed a mean number of ghost boutons of 6.5 ± 0.73 (Fig 2E; n = 10), a number not statistically different from the one observed in the stimulated preparations carrying the silenced dominant negative construct (7.75 ± 0.93; n = 12; p = 0.42) or the silenced constitutively active construct (5.9 ± 0.5; n = 10; p = 0.82). This suggests that the expression of these transgenes is efficiently suppressed by Gal80^TS^ at 18°C or that it does not affect the mean ghost bouton number. Notably, we can confirm that our constructs worked effectively because the expression of either Rho1^DN^ or Rho1^CA^ at 25°C was lethal. We therefore shifted the temperature to 29°C 48 hours before assessing ADSP. In this condition, we observed a mean number of ghost boutons of 11.14 ± 0.68 (Fig 2G; n = 35) in control animals, while animals expressing the dominant negative form of Rho1 presented a mean ghost bouton number of 12.10 ± 1.5 (n = 20), a value similar to control (Fig.2 G; p > 0.99). This suggests that Rho1 activity is not required for the formation of the morphological synaptic structures associated with ADSP. In contrast, when we expressed the constitutively active version of Rho1 in neurons 48 hours before repeated stimulation we could observe a decrease in the mean number of GB formed (Fig.2 G; 5.6 ± 0.27; n = 15; p = 0.0004). These results show that the Rho1 GTPase activity can suppress part of the morphological synaptic response that is associated with activity-dependent synaptic plasticity. This is reminiscent of its role in neuronal morphogenesis and structural plasticity in which Rho1 signaling regulates repulsive axon guidance cues and axon or dendritic retraction (Luo, 2002).

**Figure 2:**
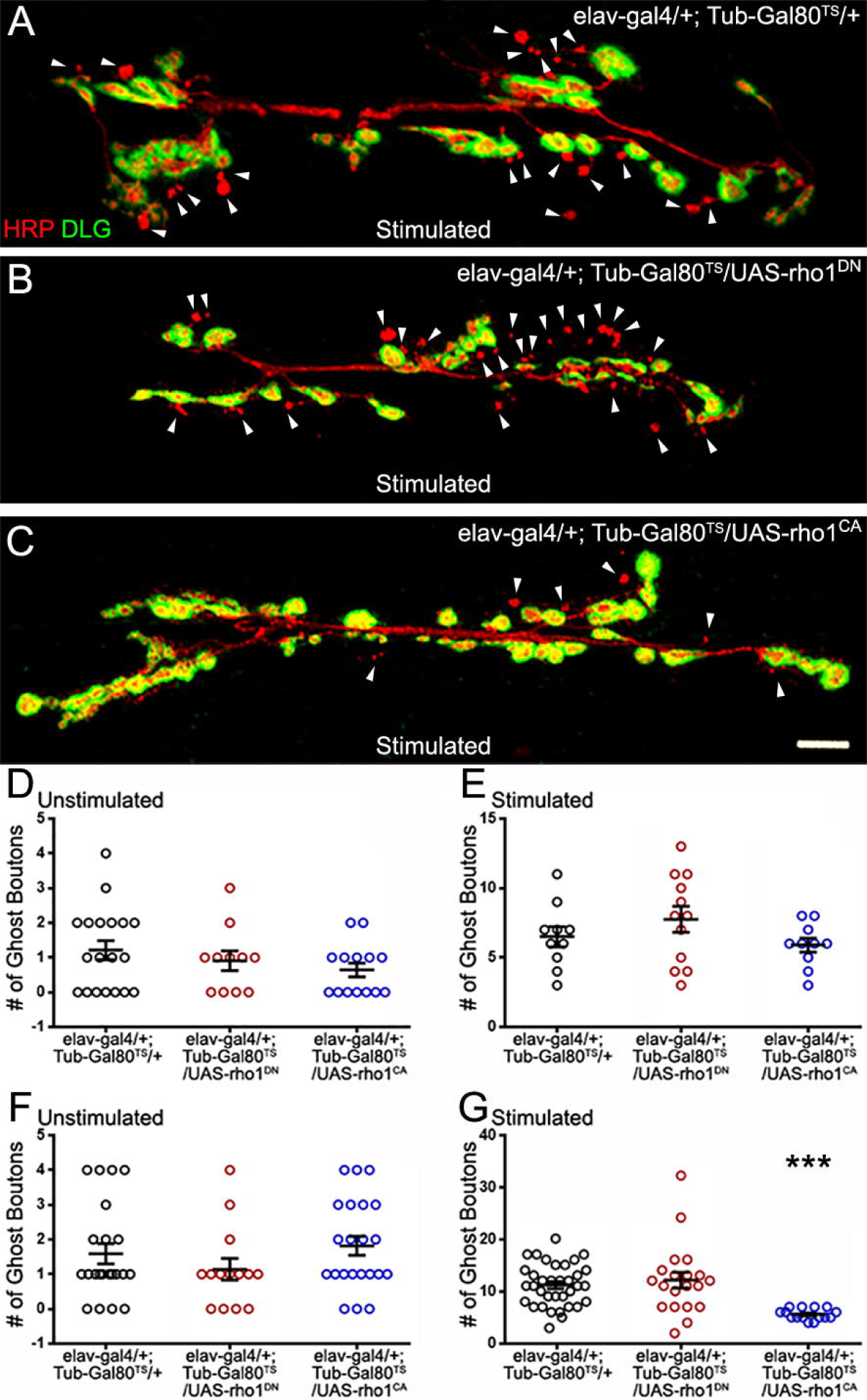
The GTPase Rho1 can repress ADSP. **(A)** Representative confocal photograph of a stimulated control synapse showing HRP and DLG immunoreactivity and ghost bouton formation (arrowheads). **(B)** Representative confocal photograph of a stimulated synapse expressing a dominant negative (DN) form of the Rho1 GTPase **(C)** Representative confocal photograph of a stimulated synapse expressing a constitutively active (CA) form of the Rho1 GTPase. **(D-G)** Quantification of mean ghost bouton number in **(D)** unstimulated and **(E)**stimulated preparations containing the repressed (18°C) conditional expression system to express Rho1^DN^ or Rho1^CA^; **(F)** unstimulated and **(G)** stimulated preparations expressing (29°C) Rho1^DN^ or Rho1^CA^ 48 hours prior to the stimulation. Scale bar: 10 µm. ANOVA with Dunnett’s multiple comparisons test: ***p < 0.001. Data are shown as scatter plots and means ± SEM.

### The GTPase Cdc42 represses ADSP

We then investigated the role of another GTPase, Cdc42, known to be activated downstream of Dsh (Moriguchi et al., 1999; Schlessinger et al., 2007). We first constitutively expressed dominant negative (DN) or constitutively active (CA) versions of Cdc42 in post-mitotic neurons (*elav-Gal4* driver). Under these conditions, we did not observe any differences in the mean number of ghost boutons in unstimulated and stimulated preparations (Fig.3 D and E). This suggests that changing Cdc42 activity might have no consequences on the neuronal properties underlying ADSP or that these consequences might be compensated for during development. To distinguish between these two possibilities, we used the previously described inducible expression system in which expression of the altered Cdc42 constructs is switched on 48 hours before carrying out the repeated stimulation protocol. As previously described, we confirmed again that the system was inhibited at 18°C (data not shown) before de-repressing the expression system 48 hours prior to the repeated stimulation protocol. Following this activation, we observed no change in unstimulated preparations (Fig. 3F). In contrast, after repeated stimulations we observed an increase in the mean number of ghost boutons when we expressed the dominant negative form of Cdc42 (compare 8.61 ± 0.46 in control to 11.23 ± 4.46 in cdc42^DN^; p = 0.009; Fig. 3B and G). Interestingly, the expression of the constitutively active form of Cdc42 showed a decrease in the number of morphological changes associated with ADSP (average of ghost bouton is 5.83 ± 0.58 in cdc42CA; p = 0.002; Fig. 3C and G). These results suggest that Cdc42 is involved in repressing the expression of ADSP.

**Figure 3:**
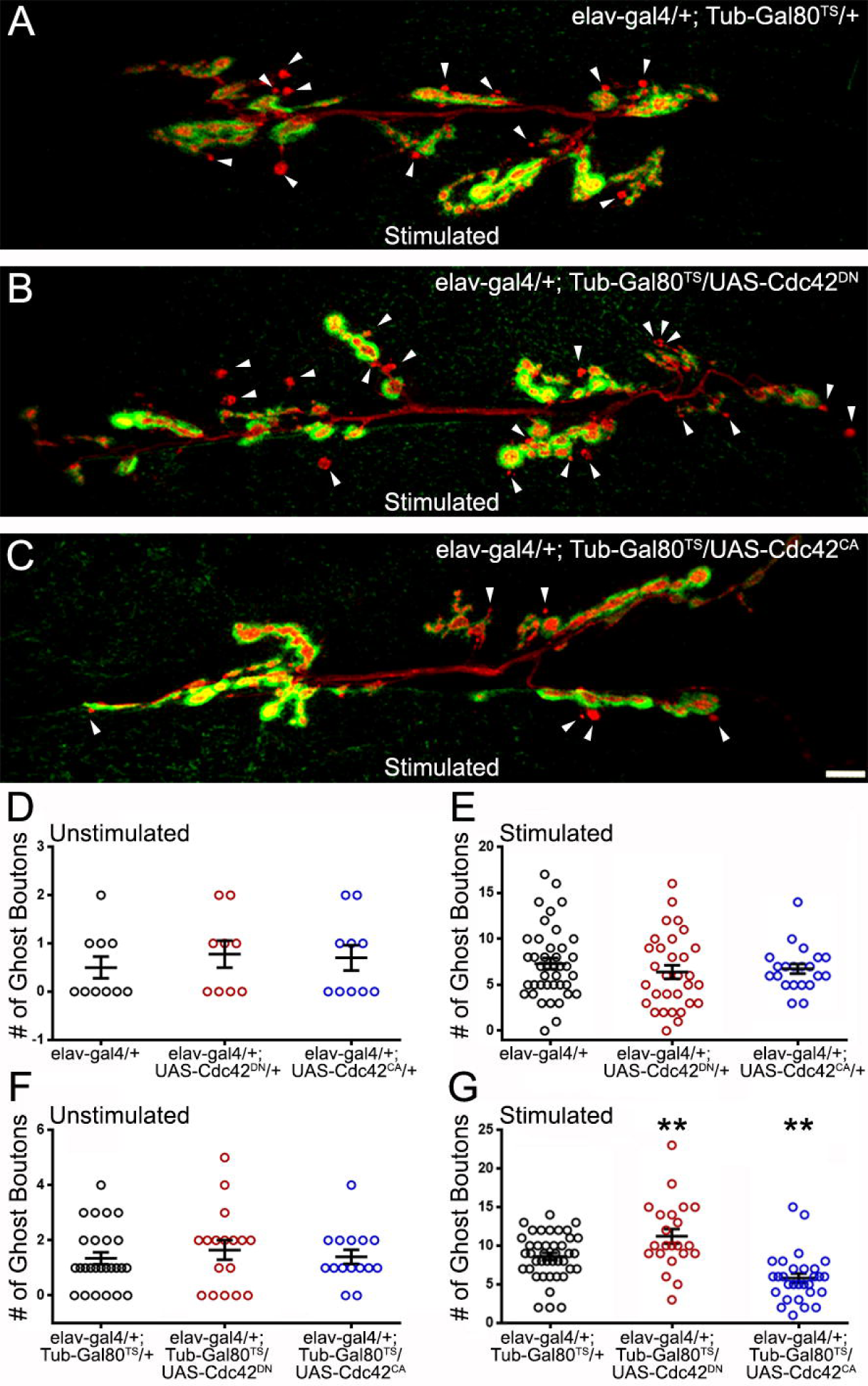
The GTPase Cdc42 represses ADSP. **(A)** Representative confocal photograph of a stimulated control synapse showing HRP and DLG immunoreactivity and ghost bouton formation (arrowheads). **(B)** Representative confocal photograph of a stimulated synapse expressing a dominant negative (DN) form of the Cdc42 GTPase **(C)** Representative confocal photograph of a stimulated synapse expressing a constitutively active (CA) form of the Cdc42 GTPase. **(D-G)** Quantification of mean ghost bouton number in **(D)** unstimulated and **(E)** stimulated control and preparations expressing Cdc42^DN^ or Cdc42^CA^ constitutively; **(F)** unstimulated and **(G)** stimulated control and preparations expressing Cdc42^DN^ or Cdc42^CA^ for 48 hours prior to the stimulation. Scale bar: 10 µm. ANOVA with Dunnett’s multiple comparisons test: **p < 0.01. Data are shown as scatter plots and mean ± SEM.

### The GTPase Rac 1 affects ADSP

We tested a third GTPase activated by Dsh and the Wg signaling, Rac1 (Bikkavilli et al., 2008; Fanto et al., 2000; Li et al., 2016), for its potential role in shaping activity-dependent synaptic plasticity. As previously described, we used a temperature-driven conditional expression system to express Rac1 dominant negative (DN) or constitutively active (CA) versions. We first verified that no phenotypes were associated with Rac1 expression at low temperatures (18°C; data not shown). We then induced Rac1^DN^ or Rac1^CA^ expression 48 hours before our experimental procedures. While we observed no differences in unstimulated preparations (Fig 4 D), we did observe changes in the mean number of ghost boutons present at the Rac1^DN^- and Rac1^CA^-expressing synapses. When control synapses showed a mean number of ghost boutons of 9.62 ± 0.42 (dark circles; n = 65; Fig.4 A), synapses expressing both Rac1 constructs showed an increase in the mean ghost bouton number. Indeed, Rac1^DN^ expressing synapses showed 12.93 ± 1.2 ghost boutons (red circles; n = 30; p = 0.002; Fig.4 B and E) and Rac1CA expressing synapses showed 14.2 ± 1.3 (blue circles; n = 18; p = 0.0009; Fig.4 C and E). Although it seems counterintuitive that the expression of both the dominant negative and the constitutively active forms of Rac1 provokes the same phenotype it is reminiscent of phenotypes previously observed during axon pathfinding within neurons expressing Rac1^DN^ or Rac1^CA^. Indeed, ISNb axons of motoneurons expressing either Rac1^DN^ or Rac1^CA^ during embryonic development showed arrested growth cones (Kaufmann et al., 1998). In addition, the expression of both Rac1^DN^ and Rac1^CA^ during embryonic axonal outgrowth causes increased axonal loss (Luo et al., 1994). Interestingly, that study showed that Rac1^CA^ expression induced a stronger phenotype and an accumulation of F-actin that was not present in Rac1^DN^-expressing animals (Luo et al., 1994), suggesting that Rac1^CA^ and Rac1^DN^ expression led to the same phenotype by affecting different cytoskeletal processes. Interestingly, at the NMJ expression of Rac1^DN^ (Park et al., 2022) and Rac1^CA^ (Ball et al., 2010) both cause an increase of synaptic boutons. Here we show that changing Rac1 activity increases the morphological changes associated with activity-dependent synaptic plasticity. Taken together, these results show that the Rho family GTPases downstream of Dsh are involved in the morphological response associated with activity-dependent synaptic plasticity.

**Figure 4:**
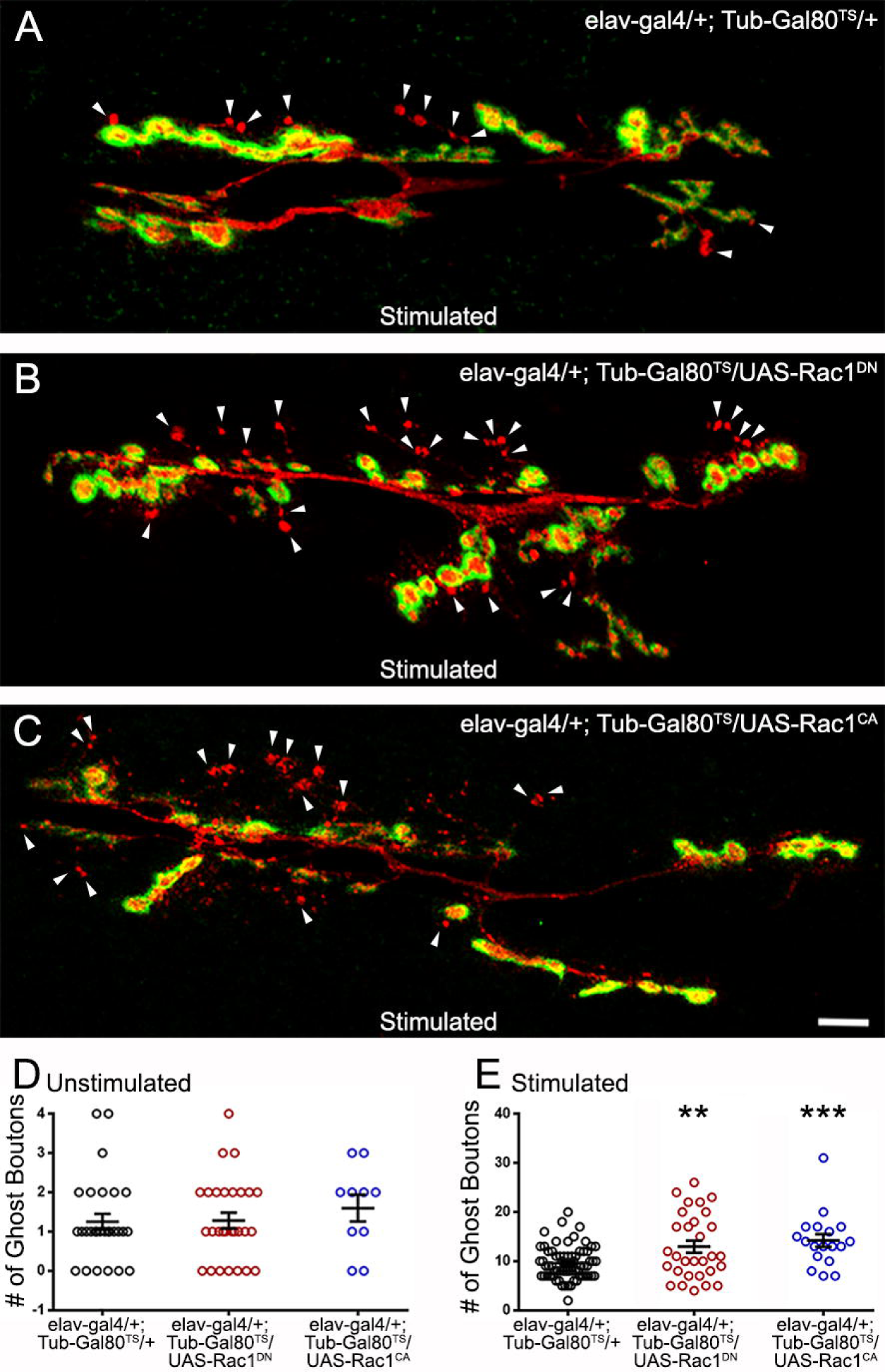
The GTPase Rac 1 can regulate ADSP. **(A)** Representative confocal photograph of a stimulated control synapse showing HRP and DLG immunoreactivity and ghost bouton formation (arrowheads). **(B)** Representative confocal photograph of a stimulated synapse expressing a dominant negative (DN) form of the Rac1 GTPase. **(C)** Representative confocal photograph of a stimulated synapse expressing a constitutively active (CA) form of the Rac1 GTPase. Quantification of mean ghost bouton number in **(D)** unstimulated and **(E)** stimulated control and preparations expressing Cdc42^DN^ or Cdc42^CA^ for 48 hours prior to the stimulation. Scale bar: 10 µm. ANOVA with Dunnett’s multiple comparisons test: **p < 0.01; ***p < 0.001. Data shown as scatter plots and mean ± SEM.

### Jun kinase activity is required for optimal ADSP in an AP1-independent manner

One of the major PCP pathway effectors is Jun kinase (Jnk; Boutros et al., 1998; Jenny, 2010; Strutt et al., 1997). This kinase can be activated by the Rho GTPases and can in turn activate cytoskeletal effectors (Bikkavilli et al., 2008; Fanto et al., 2000; Strutt et al., 1997) and transcriptional regulators. The latter are composed of the transcription factors cJun and cFos which act as dimers known as AP1 (Curran & Franza, 1988; Karin et al., 1997). We expressed a dominant-negative form of Jun kinase (Jnk^DN^) in a constitutive manner (Fig. 5E) and used the temperature-driven conditional expression (48 hours prior to repeated stimulations; Fig. 5B and G). In both cases, we observed a significant decrease in ghost bouton formation. In the experiment in which we use constitutive expression, control preparations showed a mean of 6.82 ± 0.52 (Fig. 5E) ghost boutons after stimulation, while the mean ghost bouton number of preparations with Jnk^DN^ constitutive expression was 2.47 ± 0.6 (Fig. 5E; p < 0.0001). When we use the conditional expression system, the control synapses showed 8.97 ± 0.57 (Fig. 5A and G) but only 5.63 ± 0.61 (Fig. 5B and G; p = 0.0019) in preparations presenting Jnk^DN^ conditional expression. This shows that Jnk activity is required to fully achieve the morphological changes associated with ADSP.

**Figure 5:**
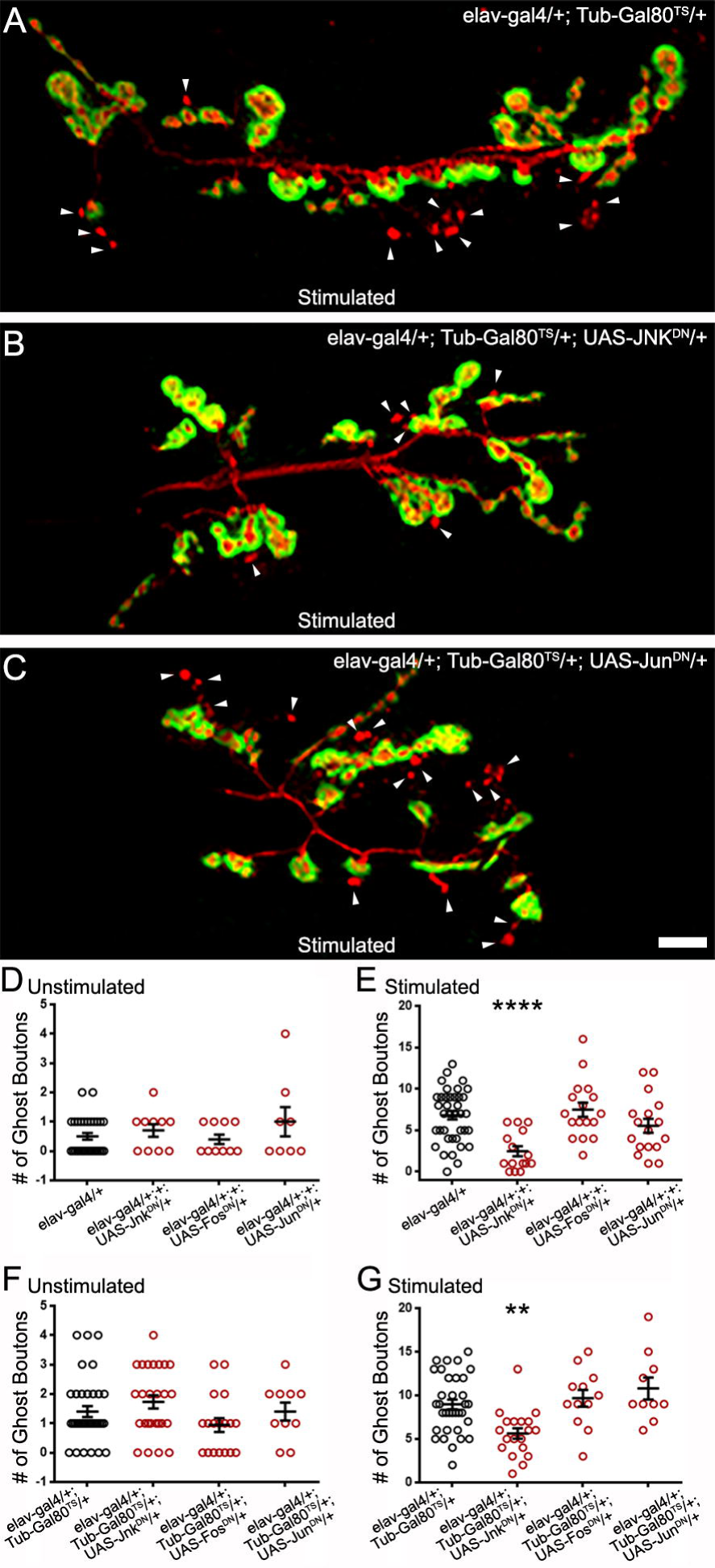
Jun Kinase is required for optimal ADSP in an AP1-independent manner. **(A)** Representative confocal photograph of a stimulated control synapse showing HRP and DLG immunoreactivity and ghost bouton formation (arrowheads). **(B)** Representative confocal photograph of a stimulated synapse expressing a dominant negative (DN) form of Jun Kinase **(C)** Representative confocal photograph of a stimulated synapse expressing a dominant negative (DN) form of c Jun. **(D-G)** Quantification of mean ghost bouton number in **(D)** unstimulated and **(E)** stimulated control and preparations constitutively expressing Jnk^DN^ or c Jun^DN^ or c Fos^DN^; **(F)** unstimulated and **(G)** preparations expressing Jnk^DN^ or c Jun^DN^ or c Fos^DN^ for 48 hours prior to the stimulation. Scale bar: 10 µm. (D) Kruskal-Wallis with Dunn’s multiple comparisons test (E-G) ANOVA with Dunnett’s multiple comparisons test: **p < 0.01; ****p < 0.0001. Data are shown as scatter plots and mean ± SEM.

We then logically asked whether the transcription factors cJun and cFos were involved in transducing the signal necessary for the ghost bouton formation after stimulation. To do so, we expressed dominant negative forms (Eresh et al., 1997) of these two constructs in a constitutive and conditional form (Fig.5 C-G). We observed no difference between control preparations and preparations containing the dominant negative forms in unstimulated and stimulated procedures. When expressing the dominant negative constructs we noticed smaller synapses, a phenotype documented before (Ballard et al., 2014) that validates our expression system. This result suggests that the transcriptional control exercised by AP1 is not necessary for the morphological changes associated with activity-dependent synaptic plasticity. We favor the hypothesis that Jnk and the Rho GTPases act on cytoskeletal regulators to achieve these morphological changes.

## Discussion

Structural synaptic plasticity is a fundamental feature of the nervous system, in which Wg/Wnt signaling plays an essential role. Reduction of Wg and its receptor Fz2 impaired the formation of activity-dependent modifications of synaptic structure (Alicea et al., 2017; Ataman et al., 2008). Overexpression of GSK3β/Sgg, an important canonical pathway member, prevents ADSP at the Drosophila NMJ (Ataman et al., 2008). Upon stimulation, Wg protein release increases at the synapse (Ataman et al., 2008), which suggests an increased activation of Wg signaling pathways. Repeated simulation of the synapse also promotes an increase of CaMKII, and reducing synaptic CaMKII impairs the activity-dependent formation of de novo boutons (Nesler et al., 2016). This suggests some role of the Wnt/Ca^2+^ non-canonical pathway on activity-dependent synaptic plasticity.

The Wg signal for polarity has been described extensively in several cellular environments (Maung & Jenny, 2011). In the developing eye, alteration of the PCP pathway provokes defects in the anterior-posterior and dorsal-ventral axes in the arrangement of ommatidia (Adler, 2002; Boutros et al., 1998; Fanto et al., 2000; Strutt et al., 1997; Strutt et al., 2013). In the Drosophila wing, the orientation of cellular hairs in the proximal-distal axis is impaired by defects of the PCP pathway (Adler, 2002; Fagan et al., 2014; Strutt et al., 1997; Wu et al., 2013). Neurons are one of the most morphologically polarized cell types, and mutations of Wnt/PCP pathway members like Dsh, Prickle, Strabismus, and Fz provoke axonal branching defects in mushroom bodies (MB) of *Drosophila* adult brain (Ng, 2012). Studies showed that Rac GTPase inactivation in mushroom bodies results in axonal branching defects (Ng et al., 2002). Wnt5, Fz, and Dsh act in concert with the small GTPase Rac1 to activate the actin assembly functions of dDAAM (dsh-associated activator of morphogenesis) necessary for the correct targeting of MB axons (Gombos et al., 2015). In addition, dDAAM regulates microtubule stability and synaptic growth at the *Drosophila* NMJ (Migh et al., 2018) while expression of Rac^DN^ (Park et al., 2022) and Rac^CA^ (Ball et al., 2010) promote synaptic overgrowth at the NMJ.

Despite this interest in the Wnt/Wg PCP pathway, its role regarding processes of synaptic plasticity has not been extensively investigated. Here, we have provided evidence that key molecules involved in the PCP pathway can affect the morphological modifications consequent to ADSP and are therefore necessary to achieve optimal activity-dependent synaptic plasticity. It is important to note that we never observed a complete ghost bouton elimination after affecting the PCP pathway. This makes sense since the other Wg pathways — the canonical pathway (Alicea et al., 2017; Ataman et al., 2008) and the calcium pathway (Freeman et al., 2011; Nesler et al., 2016)— are also necessary to achieve full activity-dependent synaptic plasticity. The contribution of the Wg PCP pathway to the formation of *de novo* synaptic structures after stimulation illustrates the redundancy and robustness of the Wg/Wnt signal in directing activity-dependent synaptic plasticity.

Our work revealed PCP pathway molecules exercising both activation and repression of activity-dependent synaptic plasticity. Indeed, the loss of function of *dsh*, either using RNAi or the PCP-specific allele *dsh^1^*, showed compromised plasticity. The same was true for the loss of function of *jnk*. This suggests that the role of these molecules is to promote activity. Activation and repression of ghost bouton formation was also achieved by modifying Rho GTPase activity. Expression of the constitutively active forms of the three GTPases we tested showed phenotypes, suggesting that these molecules can regulate ADSP. While Cdc42^CA^ and Rho1^CA^ expressing synapses show reduced plasticity, Rac1^CA^ expression elicited an over plastic response. This antagonistic relationship between Rho1 and Rac1 has been documented previously. For example, Rac1 activation promotes neurite formation and outgrowth, but Rho1 activation leads to neurite retraction and suppresses neurite outgrowth (Lorenzetto et al., 2013; Luo, 2000). Similarly, proper dendritic spine morphogenesis requires balanced Rho GTPase activity regulation (Penzes & Cahill, 2012). Interestingly, Rac1 induces the formation and maintenance of dendritic spines, while Rho1 decreases spine formation by promoting spine pruning or retraction (Bolognin et al., 2014; Nakayama et al., 2000; Newey et al., 2005; Tashiro et al., 2000).

The observed Cdc42 phenotypes indicate a role in repressing activity-dependent synaptic plasticity. Indeed, the dominant negative form is over plastic, while the constitutively active form is under plastic. This shows that in our system Cdc42 represses ghost bouton formation. This is surprising since most studies show that Cdc42 and the Rho GTPases in general promote membrane protrusion (Sadok & Marshall, 2014). For example, Rac1 and Cdc42 both promote dendritic spine formation (Bolognin et al., 2014; Nakayama & Luo, 2000), cell extension (Nakayama et al., 2000; Swetman et al., 2002), and actin polymerization (Lamarche et al., 1996; Nobes & Hall, 1995). At the NMJ, *cdc42* mutations provoke an overgrown synapse, suggesting that Cdc42 represses synaptic growth (Rodal et al., 2008) while expression of Rac^DN^ (Park et al., 2022), Rac^CA^ and Rac^OE^ (Ball et al., 2010) promotes synaptic growth. These studies might indicate a more complex interaction between the Rho GTPases.

While Rho1^CA^ animals showed repressed activity-dependent synaptic plasticity, Rho1^DN^ did not show any phenotype. It is important to note that we know that Rho1^DN^ expression can decrease Rho function since its constitutive expression creates lethality. We can therefore hypothesize that the absence of phenotype in the Rho1^DN^ animals might be due to compensation/redundancy from other GTPase activity. Alternatively, its function at the time at which we express its dominant negative form is perhaps not to repress activity-dependent synaptic plasticity. Indeed, Rho GTPases serve different functions at different times. For example, in hippocampal neurogenesis, Cdc42 promotes initial dendritic development and spine maturation, while Rac1 is essential for the late stages of dendritic growth and spine maturation (Vadodaria et al., 2013).

Finally, both the constitutive and dominant negative forms of Rac1 provoke over plasticity at the NMJ. Although surprising, this is not the first time that the same phenotype has been observed when expressing these two constructs. Expression of both constitutively active and dominant-negative forms of Rac1 causes axonal loss in Drosophila (Luo et al., 1994), and Rac1 can have a dual function regulating attractive and repulsive signals in axonal guidance (Fan et al., 2003). Taken together, these results give an image of an intricate and complex local role of the Rho GTPases in regulating ADSP induced at the synapse upon high neuronal activity.

We have shown that the transcriptional regulator AP1 is unlikely to be involved in activity-dependent synaptic plasticity. We favor the hypothesis that regulation of the cytoskeleton by the PCP pathway is involved in regulating this plasticity. The reorganization of the synaptic cytoskeleton has been extensively studied in spines and is linked to processes of activity-dependent plasticity and learning and memory (Cornelius et al., 2021; Macgillavry et al., 2016; Repetto et al., 2014). This process involves the reorganization of filamentous actin, actin regulators, and microtubules (Cornelius et al., 2021; Ka & Kim, 2016; Schätzle et al., 2018). In fact, we have previously isolated one such regulator, the actin regulator Cortactin (Weaver et al., 2001; Weed et al., 2000) and showed that it is a regulator of activity-dependent synaptic plasticity, and that its protein is upregulated under the control of Wg signaling (Alicea et al., 2017). Determining whether the PCP pathway is involved in this regulation will be a challenge for the future.

## Author Contributions

CDC: Conceptualization, Formal analysis, Investigation, Visualization, Writing – original draft; MV: Investigation, Writing – review & editing; BM: Conceptualization, Funding acquisition, Supervision, Visualization, Writing – original draft

## Acknowledgments

We thank Dr. Jonathan Blagburn for his valuable comments on previous versions of this manuscript.

Parts of our schematic diagrams (Figure 1 A and E) were created with Biorender.com.

